# A neuronal mechanism underlying decision-making deficits during hyperdopaminergic states

**DOI:** 10.1101/211862

**Authors:** Jeroen P.H. Verharen, Johannes W. de Jong, Theresia J.M. Roelofs, Christiaan F.M. Huffels, Ruud van Zessen, Mieneke C.M. Luijendijk, Ralph Hamelink, Ingo Willuhn, Hanneke E.M. den Ouden, Geoffrey van der Plasse, Roger A.H. Adan, Louk J.M.J. Vanderschuren

**Author notes:** These authors contributed equally (Roger A.H. Adan, Louk J.M.J. Vanderschuren). Correspondence should be addressed to R.A.H.A. or L.J.M.J.V.

## Abstract

Hyperdopaminergic states in mental disorders are associated with disruptive deficits in decision-making. However, the precise contribution of topographically distinct mesencephalic dopamine pathways to decision-making processes remains elusive. Here we show, using a multidisciplinary approach, how hyperactivity of ascending projections from the ventral tegmental area (VTA) contributes to faulty decision-making in rats. Activation of the VTA-nucleus accumbens pathway leads to insensitivity to loss and punishment due to impaired processing of negative reward prediction errors. In contrast, activation of the VTA-prefrontal cortex pathway promotes risky decision-making without affecting the ability to choose the economically most beneficial option. Together, these findings show how malfunction of ascending VTA projections affects value-based decision-making, providing a mechanistic understanding of the reckless behaviors seen in substance abuse, mania, and after dopamine replacement therapy in Parkinson’s disease.

---

Impaired decision-making can have profound negative consequences, both in the short and in the long term. As such, it is observed in a variety of mental disorders, such as mania^1,2^, substance addiction^3–6^, and as a side effect of dopamine (DA) replacement therapy in Parkinson’s disease^7,8^. Importantly, these disorders are associated with aberrations in DAergic neurotransmission^9,10^, and DA has been implicated in decision-making processes^11–13^. However, ascending DAergic projections from the ventral mesencephalon are anatomically and functionally heterogeneous^14–16^ and the contribution of these distinct DA pathways to decision-making processes remains elusive.

The mesocorticolimbic system, comprising DA cells within the ventral tegmental area (VTA) that mainly project to the nucleus accumbens (NAc; mesoaccumbens pathway) and medial prefrontal cortex (mPFC; mesocortical pathway), has an important role in value-based learning and decision-making^14–16^. When an experienced reward is better than expected, the firing of VTA DA neurons increases, thereby signaling a discrepancy between anticipated and experienced reward to downstream regions. Conversely, when a reward does not fulfill expectations, DA neuronal activity decreases. This pattern of DA cell activity is the basis of reward prediction error (RPE) theory^17–20^, which describes an essential mechanism through which organisms learn to flexibly alter their behavior when the costs and benefits associated with different courses of action shift. Although the relevance of RPEs in value-based learning is widely acknowledged, little is known about how different VTA target regions process these DA-mediated error signals, and how this ultimately leads to adaptations in behavior.

Here, we used projection-specific chemogenetics combined with behavioral tasks, pharmacological interventions, computational modelling, *in vivo* microdialysis and *in vivo* neuronal population recordings to investigate how different ascending VTA projections contribute to value-based decision-making processes in the rat. Specifically, we investigated the mechanism underlying the aberrant decision-making style that is associated with increased DA neuron activity. We hypothesized that hyperactivation of VTA neurons interferes with reward prediction error processing, leading to impaired adaptation to reward value dynamics. We predicted an important contribution of the mesoaccumbens pathway in incorporating experienced reward, loss and punishment into future decisions, considering the importance of the NAc in reinforcement learning and motivated behaviors^21–23^, and a modulatory role for the mesocortical pathway in value-based choice behavior, given its involvement in executive functions, such as decision-making and behavioral flexibility^24,25^. Furthermore, we tested an explicit prediction based on a neurocomputational model of the DA system, in which impaired negative RPE processing is involved in learning deficits during DA replacement therapy^7,26^.

## RESULTS

### Dopaminomimetic drugs impair serial reversal learning

To test the role of DA in flexible value-based decision-making, rats were tested in a serial reversal learning task following systemic treatment with the DA neurotransmission enhancers cocaine and D-amphetamine. A reversal learning session (Fig. 1a) comprised 150 trials, and started with the illumination of two nose poke holes in an operant conditioning chamber. One of these was randomly assigned as active, and responding in this hole resulted in sucrose delivery under a fixed-ratio (FR) 1 schedule of reinforcement. When animals had made five consecutive correct responses, the contingencies reversed so that the previously inactive hole now became active, and vice versa.

**Figure 1.**
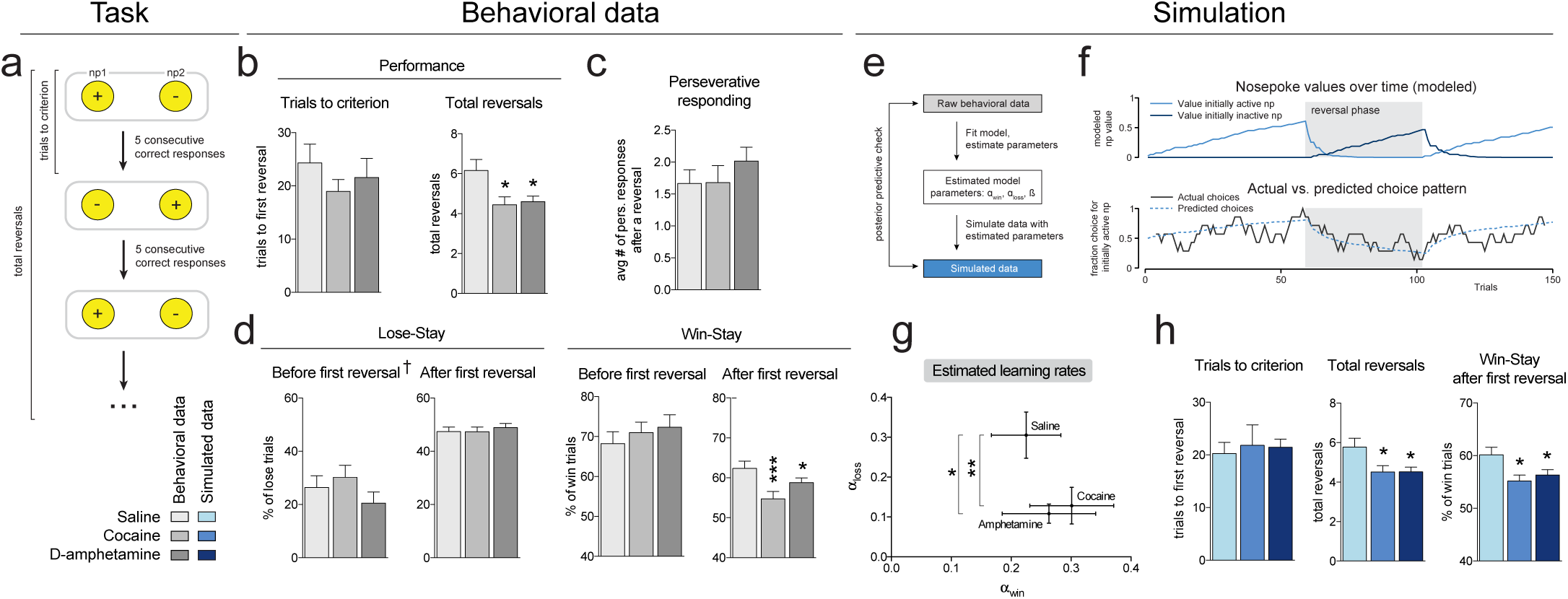
Treatment with cocaine or D-amphetamine impairs reversal learning. (**a**) Task design. (**b**) Systemic treatment with cocaine (10 mg/kg) or D-amphetamine (0.25 mg/kg) did not alter the number of trials required to reach the first reversal (one-way repeated measures ANOVA, p = 0.55). However, cocaine-or D-amphetamine treatment decreased the total number of reversals accomplished (one-way repeated measures ANOVA, p = 0.0037; post-hoc Sidak’s multiple comparisons test, p = 0.0102 cocaine versus saline, p = 0.0197 D-amphetamine versus saline). (**c**) Treatment with cocaine or D-amphetamine did not alter perseverative behavior after a reversal (p = 0.46). (**d**) Lose-stay behavior was unaffected after both cocaine or D-amphetamine treatment, both before (p = 0.21 ^†^) and after (p = 0.77) the first reversal. Cocaine and D-amphetamine decreased win-stay behavior after (ANOVA, p = 0.0007; post-hoc Sidak’s multiple comparisons test, p = 0.0009 for cocaine versus saline, p = 0.0336, D-amphetamine versus saline), but not before the first reversal (p = 0.67). Data in (b),(c),(d) and (g): repeated measures from n = 25 animals. ^†^ 6 animals had no losses before the first reversal (i.e., trials to first criterion was 5), so the repeated measures ANOVA was performed on data of n = 19 animals; graph shows n = 25. (**e**) We used a modified Rescorla-Wagner model to describe the behavior of the rats during reversal learning. (**f**) Simulated data from an example session. (upper panel) Simulated values of the nose pokes, given the rat’s optimal model parameters and observed choice sequence. (lower panel) Modeled choice probabilities, converted from the simulated nosepoke values using a softmax (unsmoothed), and the rat’s actual choice pattern (smoothed over 7 trials). (**g**) Best-fit learning parameters. Treatment with cocaine and D-amphetamine significantly decreased α_loss_, without affecting the other model coefficients. (Wilcoxon matched pairs signed rank test, * p = 0.032, ** p = 0.0046, see also Table S2) (**h**) Simulating data with the model parameters extracted in (g) replicated the drug-induced effects of the behavioral data shown in (b) and (d). (n = 25 simulated rats; ANOVA on trials to criterion, p = 0.86; ANOVA on total reversals, p = 0.0114, post-hoc Sidak’s test, p = 0.0411 for cocaine and p = 0.0215 for D-amphetamine; ANOVA on win-stay behavior, p = 0.0090, post-hoc Sidak’s test, p = 0.0181 for cocaine and p = 0.0462 for D-amphetamine. ANOVA on all other outcomes measures, all p > 0.1). Data are shown as mean ± standard error of the mean.

Injection of either drug did not affect the number of trials needed to reach the criterion of a series of five consecutive correct responses (Fig. 1b, left panel). However, the number of reversals achieved in the entire session was significantly reduced in the drug-treated animals (Fig. 1b, right panel, and Fig. S1a). Thus, cocaine and D-amphetamine impaired task performance, but this effect did not appear until the moment of first reversal. We reasoned that this pre-and post-reversal segregation in drug effects on task performance is related to the structure of the task (Fig. 1a). That is, after every reversal, the value of the outcome of responding in the previously active hole declines, and conversely, the value associated with responding in the previously inactive hole increases. Accordingly, this task entails a combination of devaluation and revaluation mechanisms following reversals.

To understand the nature of the drug-induced deficit in reversal learning performance, we analyzed the animals’ behavior in more detail. Perseverative responding, i.e. the average number of responses in the previously active hole directly after a reversal, was not altered after cocaine or D-amphetamine treatment (Fig. 1c). Lose-stay behavior, i.e. the percentage of (unrewarded) trials in the inactive nose poke hole followed by a response in the (still) inactive hole, was also not affected (Fig. 1d, left panel). However, win-stay behavior, i.e. the percentage of responses in the active nose poke hole after which the animal responded in that same active hole, was significantly decreased after treatment with cocaine or D-amphetamine (Fig. 1d, right panel). This drug-induced reduction in win-stay behavior indicates that even though the animals received a reward after responding in the active nose poke hole, they next sampled the inactive hole more often than after saline treatment. Importantly, win-stay behavior was only reduced after reversal, indicating that behavioral impairments were not the result of a general decline in task performance or sensitivity to reward.

Overall, the effects in the reversal learning task indicate that increased DA signaling after cocaine or D-amphetamine treatment did not impair the animals’ ability to find the active nose poke hole at task initiation, hence to assign positive value to an action. Yet, when the values of (the outcome of) two similar actions (that is, responding in a nose poke hole) changed relative to each other, drug-treated animals were impaired in adjusting behavior, perhaps as a result of a valuation deficit. This suggests that treatment with these drugs disrupted the process of integrating recent wins or losses (i.e., a revaluation or a devaluation impairment, respectively) in decisions.

To gain insight into the mechanisms underlying impaired reversal learning, we modelled the behavior of each subject by fitting the data to a computational reinforcement learning model (Fig. 1e,f and Table S1). We used an extended version of the Rescorla-Wagner model^27,28^, using two different learning rates, ɑ_win_ and ɑ_loss_, describing the animal’s ability to learn from wins and losses, respectively^29^. Such a model-based approach investigates task performance based on an extended history of trial outcomes, and not merely the most recent outcome, such as win-and lose-stay measures do, providing a more in-depth analysis of the learning capacity of the animals.

When comparing the Rescorla-Wagner model coefficients of the animals after saline with those after cocaine and D-amphetamine treatment, we observed a strong decrease in parameter ɑ_loss_ without affecting ɑ_win_ or choice stochasticity factor β (Fig. 1g,h, Fig. S1b,c and Table S2). This indicates that cocaine and D-amphetamine interfere with learning from negative, but not positive, RPEs.

### Chemogenetic activation of mesoaccumbens pathway impairs reversal learning

In view of the role of DA in RPE signaling, we hypothesized that cocaine and D-amphetamine interfered with learning from losses by overactivation of ascending midbrain DA projections, thereby disrupting negative RPEs. This same mechanism has been hypothesized to be involved in the DA dysregulation syndrome in medicated Parkinson’s disease patients^7,30^. Such an overactivation may lead to an inability to devalue stimuli and/or their associated outcomes, resulting in choice behavior that is not optimally value-based. Specifically, we were interested in the contribution of projections from the VTA to the NAc and the mPFC to impairments in reversal learning.

In order to activate neuronal subpopulations of the VTA in a projection-specific manner, we combined a canine adeno-associated virus retrogradely delivering Cre-recombinase (CAV2-Cre) and a Cre-dependent viral vector encoding hM3Dq(Gq)-DREADD fused to mCherry-fluorescent protein^31^ (Fig. 2a and Fig. S2). This two-viral approach resulted in high levels of DA specificity (80% of the transfected neurons in the mesoaccumbens group and 72% of the transfected neurons in the mesocortical group were positive for tyrosine hydroxylase, Fig. 2b). To investigate whether the effects of cocaine and D-amphetamine on reversal learning were driven by activation of the mesoaccumbens or mesocortical pathway, animals were injected with clozapine-N-oxide (CNO) immediately before testing in the reversal learning task.

**Figure 2.**
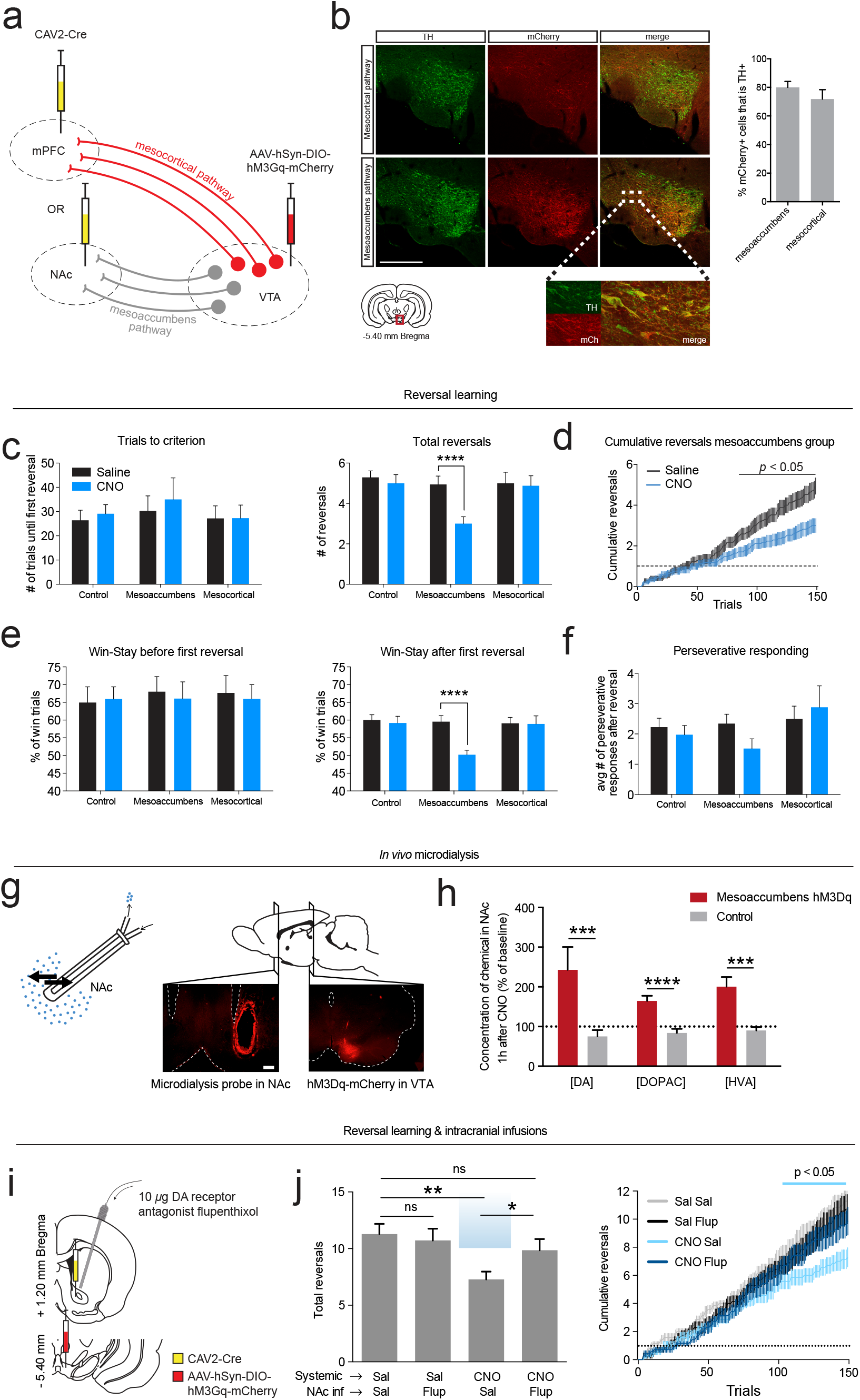
Chemogenetic activation of the mesoaccumbens, but not mesocortical pathway mimicked the effects of cocaine and D-amphetamine on reversal learning. (**a**) Experimental design. Animals received an infusion of CAV2-Cre into either the mPFC or NAc. A Cre-dependent Gq-DREADD virus was injected bilaterally into the VTA. (**b**) (left panel) Representative histology images showing coronal sections stained for tyrosine hydroxylase (left), DREADD-mCherry (middle) and an overlay (right). Image bottom left corner from Paxinos and Watson (2007). Scalebar, 500 μm. (right panel) Co-staining of mCherry with tyrosine hydroxylase, showing the percentage of DREADD-transfected neurons that is dopaminergic (mean ± s.d.). Data from n = 9 (mesoaccumbens), n = 8 (mesocortical) animals. (**c**) (left panel) Activation of either pathway did not affect the number of trials needed to reach the first reversal (i.e., 5 consecutive correct responses; two-way repeated measures ANOVA; main effect of CNO, p = 0.54; group × CNO interaction, p = 0.90). (right panel) Performance on the task over the entire session was significantly impaired after mesoaccumbens activation (two-way repeated measures ANOVA; main effect of CNO, p = 0.0025; group × CNO interaction, p = 0.0067; post-hoc Sidak’s multiple comparisons test, p = 0.89 for control group, p < 0.0001 for mesoaccumbens group, p = 0.99 for mesocortical group) (**d**) Plot of the cumulative reversals over time shows that the performance deficit after mesoaccumbens activation does not appear until after the first reversal (Sidak’s multiple comparisons test corrected for 150 comparisons, p < 0.05 after trial 85). Dashed line indicates first reversal. (**e**) A significant decrease in win-stay behavior after (two-way repeated measures ANOVA; main effect of CNO, p = 0.0040; group × CNO interaction, p = 0.0026; post-hoc Sidak’s multiple comparisons test, p = 0.9647 for control group, p < 0.0001 for mesoaccumbens group, p = 0.9997 for mesocortical group), but not before first reversal (two-way repeated measures ANOVA; main effect of CNO, p = 0.78; group × CNO interaction, p = 0.91) was observed during mesoaccumbens activation. (**f**) Perseverative behavior was not affected (two-way repeated measures ANOVA; main effect of CNO, p = 0.89; group × CNO interaction, p = 0.71). All data: n = 17 control, n = 17 mesoaccumbens, n = 16 mesocortical group. (**g**) Microdialysis was used to measure extracellular concentrations of DA and its metabolites in the NAc after chemogenetic mesoaccumbens stimulation. Scalebar, 500 μm. (**h**) NAc levels of DA and its metabolites were elevated one hour after an i.p. CNO injection in DREADD-infected animals compared to controls (post-hoc tests, DA, p = 0.0002; DOPAC, p < 0.0001; HVA, p = 0.0008; see also Fig. S4). (**i**) Prior to reversal learning, animals received systemic CNO (or saline) for DREADD stimulation and a microinjection with α-flupenthixol (or saline) into the nucleus accumbens. (**j**) α-flupenthixol itself had no effect on reversal learning, but prevented the CNO-induced impairment on reversal learning (ANOVA, p = 0.0024; post-hoc Holm-Sidak’s test: **p = 0.0019, *p = 0.0397). Note that animals had a higher baseline of reversals in this experiment, because the animals were trained on the task (see Online methods). Abbreviations: Sal, saline; Flup, α-flupenthixol; ns, not significant.

Chemogenetic activation of the mesoaccumbens pathway resulted in the same pattern of impairments in reversal learning as cocaine and D-amphetamine treatment, i.e. a reduction in the numbers of reversals achieved, without affecting trials to first reversal criterion (Fig. 2c). This pattern was confirmed by plotting the cumulative reversals as a function of completed trials (Fig. 2d and Fig. S3a). Similar to cocaine and D-amphetamine, the performance impairment during mesoaccumbens activation was associated with a post-reversal (but not pre-reversal) decrease in win-stay behavior (Fig. 2e), whereas perseverative responding and lose-stay behavior were not altered (Fig. 2f and Fig. S3b). Remarkably, during mesoaccumbens activation, both win-and lose-stay behavior were around 50% post-reversal, indicative of random choice behavior. Indeed, the Rescorla-Wagner model fitted with a significantly lower likelihood after mesoaccumbens activation (Fig. S3c), indicating that the animals’ performance declined such that the model was less able to describe the data compared to baseline conditions. In contrast to mesoaccumbens activation, mesocortical activation or CNO injection in a sham-operated control group had no effect on reversal learning.

The finding that hyperactivity in the mesoaccumbens pathway evoked similar effects on reversal learning as cocaine and D-amphetamine did, suggests that these drugs exert their influence on flexible value-based decision-making through DA neurotransmission within the NAc. To directly test this, we performed *in vivo* microdialysis in the NAc of animals that expressed Gq-DREADD in the mesoaccumbens pathway (Fig. 2g). Administration of CNO increased baseline levels of DA in the NAc, as well as its metabolites 3,4-dihydroxyphenylacetic acid (DOPAC) and homovanillic acid (HVA) (Fig. 2h and Fig. S4). Next, we infused the DA receptor antagonist α-flupenthixol into the NAc of DREADD-treated animals prior to chemogenetic activation of the mesoaccumbens pathway in a reversal learning test (Fig. 2i). This dose of α-flupenthixol had no effect on reversal learning after systemic saline injection, but it restored the effect of chemogenetic activation of the mesoaccumbens pathway to a level statistically indistinguishable from saline treatment (Fig. 2j). This finding supports the assumption that the effects of mesoaccumbens hyperactivity are mediated through NAc DA receptor stimulation.

### Dopamine neuron activity during reversal learning

Considering the function of RPEs in value updating^20^, we tested whether midbrain DA neurons tracked the presence of wins and losses in the form of RPEs during reversal learning. To this aim, we measured *in vivo* neuronal population activity from DA neurons in the VTA using fiber photometry^32^ in TH::Cre rats (Fig. 3a and Supplementary Movie 1).

**Figure 3.**
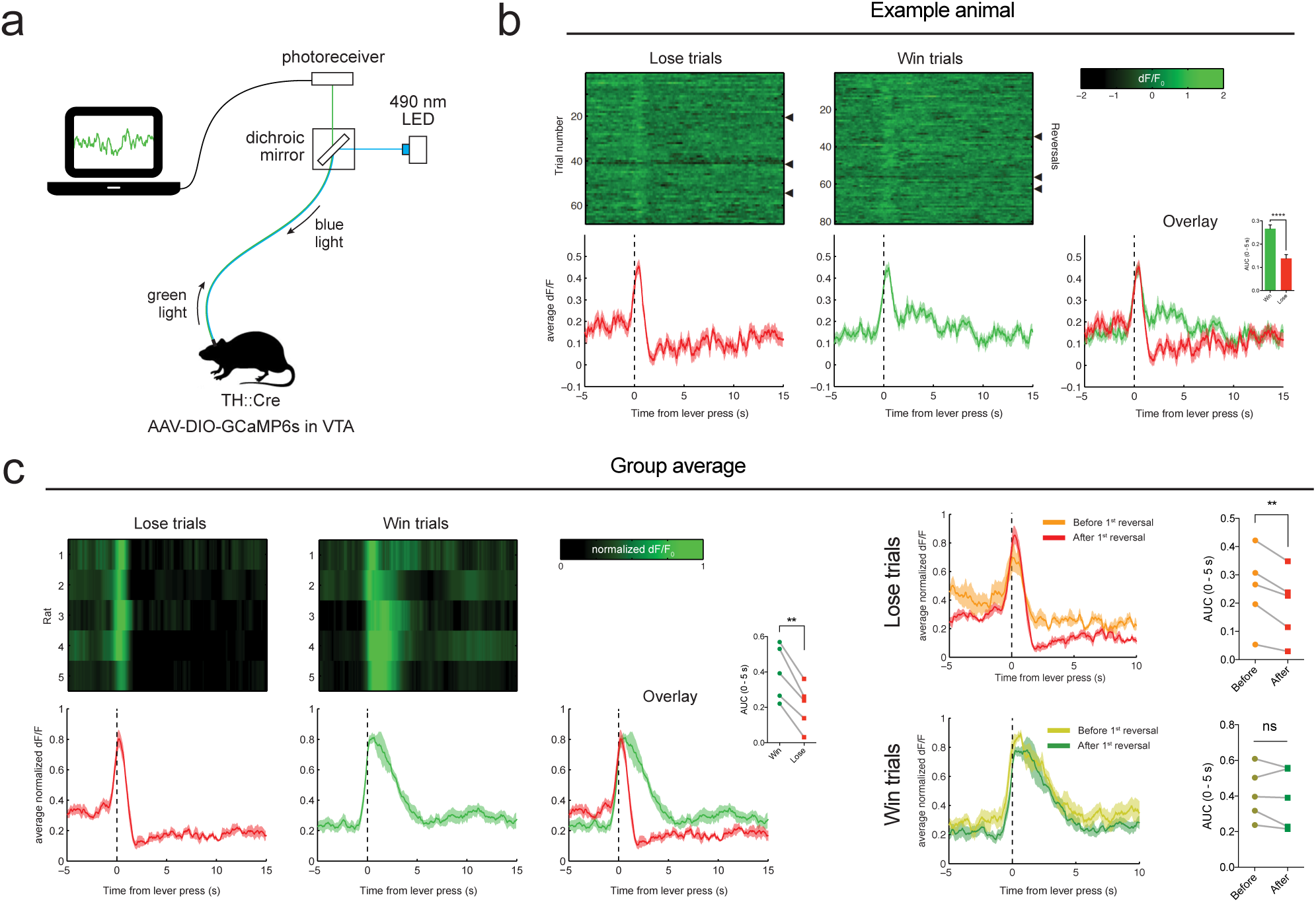
*In vivo* fiber photometry in VTA DA neurons during reversal learning. (**a**) Experimental setup. (**b**) Reversal learning session of an example animal. Triangles depict a reversal. Data is time-locked to a lever press by the rat and (in win trials) immediate reward delivery. Inset shows area under the curve in the first 5 seconds following lever press (unpaired t-test, p < 0.0001). (**c**) Group average. (left panels) VTA DA neurons responded differentially to wins and losses (AUC (inset), paired t-test, p = 0.0015). (right panels) Lose trials evoked a stronger negative reward prediction error signal after the first reversal compared to before reversal. (AUC (inset), paired t-test, p = 0.0062 for lose trials, p = 0.3658 for win trials)

Around the time of responding, we observed a clear two-component RPE signal^20^ (Fig. 3b,c and Fig. S5), i.e. a ramping of DA activity towards the moment of response, followed by an additional value component. That is, win trials were associated with a prolonged DA peak, whereas loss trials were characterized by a rapid decline in DA population activity after the response was made. No such signals were observed in animals injected with an activity-independent control fluorophore (Fig. S5).

Since mesoaccumbens hyperactivity only affected task performance after reversal, we compared DA activity pre-and post-reversal (Fig. 3c, right panels). In loss trials, we observed significantly stronger negative RPEs after the first reversal compared to before reversal. In contrast, DA peaks during the win trials were similar before and after the first reversal. This supports our notion that the impairment in reversal learning during mesoaccumbens hyperactivity was due to selective interference with learning from negative RPE-guided feedback.

### Mesoaccumbal activation interferes with adapting to devaluations

To examine whether the effects of mesoaccumbens hyperactivity on learning from negative feedback generalizes to conditions beyond reversal learning, we trained rats on a probabilistic discounting task (modified from refs. 33 and 34). In this task, rats could choose between responding on a ‘safe’ lever, which always produces one sucrose pellet, or on another, ‘risky’ lever, which produces a larger reward (i.e., three sucrose pellets) with a given probability. Within a session, the chance of receiving the large reward after a response on the risky lever decreases across four trial blocks — in the first block, animals always received the large reward when pressing the risky lever, whereas the odds of winning were reduced to 1 in 12 in the fourth block (Fig. 4a and Fig. S6a). An important difference with reversal learning is that in this task, a response shift is not the best option after a loss *per se* — lose-stay behavior at the risky lever may yield the same amount of sucrose as a shift to the safe lever, depending on the odds in the trials block. Therefore, an increase in lose-stay or decrease in win-stay behavior does not necessarily reflect poor choice behavior.

**Figure 4.**
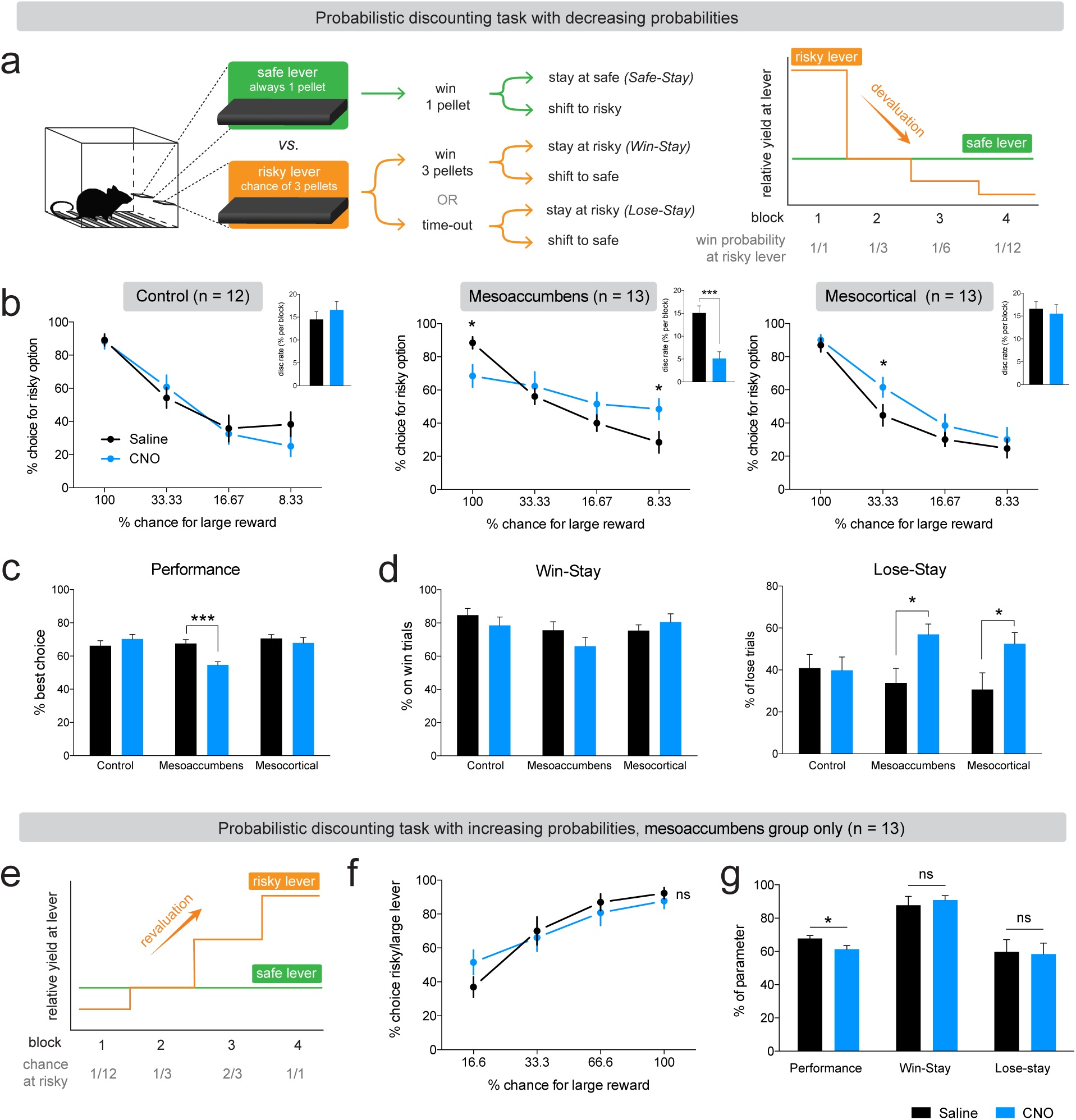
Chemogenetic activation of the mesoaccumbens and the mesocortical pathway alters probabilistic discounting. (**a**) Task design. (**b**) Discounting curves for individual groups. (left panel) Sham control group (saline vs CNO; Sidak’s test, p > 0.1 for all blocks). (middle panel) During mesoaccumbal hyperactivity, animals have a smaller preference for the risky lever in the first block (Sidak’s test, p = 0.0468), a larger preference for the risky lever in the last block (p = 0.0468; block 2 and 3 both p > 0.1), and a significantly diminished discounting rate (inset, p = 0.0002). (right panel). Mesocortical activation increased choice for the risky lever in the second block (Sidak’s test in block 2, p = 0.0247; block 1, 3 and 4, all p > 0.1). Asterisks in discounting curves indicate significant difference between saline and CNO treatment. Insets display the average steepness of the discounting curve (statistical comparison with Sidak’s test). (**c**) Mesoaccumbens activation reduces the percentage optimal choices in the probabilistic discounting task (i.e., % best choice in blocks 1, 3 and 4; two-way repeated measures ANOVA; main effect of CNO, p = 0.0331; group × CNO interaction effect, p = 0.0016; post-hoc Sidak’s test, p = 0.5082 for control group, p = 0.0004 for mesoaccumbens group, p = 0.7533 for mesocortical group). (**d**) Chemogenetic activation of the mesoaccumbens or mesocortical pathway had no effect on win-stay behavior (two-way repeated measures ANOVA; main effect of CNO, p = 0.36; group × CNO interaction effect, p = 0.26), but did increase lose-stay behavior (two-way repeated measures ANOVA; main effect of CNO, p = 0.0026; group × CNO interaction effect, p = 0.0622; post-hoc Sidak’s test, p = 0.9988, p = 0.0177 and p = 0.0203 for control, mesoaccumbens and mesocortical groups, respectively). (**e**) Task design of the probabilistic discounting task with increasing probabilities. (**f**) Mesoaccumbens activation did not affect the discounting curve (Sidak’s test in every block, p > 0.1). (**g**) Mesoaccumbens activation decreased performance on the task (paired t-test, p = 0.0143), but not win-stay (paired t-test, p = 0.32) or lose-stay behavior (paired t-test, p = 0.85). Data are shown as mean ± standard error of the mean.

After training, the animals showed stable discounting performance, preferring the risky lever in the first block, and shifting their choice towards the safe lever when the yield of the risky lever diminished (Fig. 4b, left panel). Mesoaccumbens activation (Fig. 4b, middle panel) decreased the choice of the risky lever in the first block and increased choice for the risky lever in the last block, resulting in a significantly reduced slope of the discounting curve (Fig. 4b, middle panel, inset), and a lower percentage of optimal choices (Fig. 4c). Importantly, the inability to discount the value of the risky lever in the latter blocks of the task is indicative of an inability to adapt to a declining outcome of responding on the risky lever (Fig. S6b). The reduced choice for the risky lever in the first block may also be due to a devaluation deficit, as the receipt of only one sucrose pellet after responding on the safe lever (compared to the three pellet yield of responding on the risky lever) may be perceived as a ‘loss’, since the relative value of responding on the safe lever is lower in this block^35^. In contrast, mesocortical activation only increased risk-seeking in the second block, in which the yield of the safe (1 pellet) and risky (1 in 3 chance of 3 pellets) levers were equal (Fig. 4b, right panel), so that the amount of optimal choices remained unaffected (Fig. 4c). Further analysis of task strategy showed that lose-stay behavior at the risky lever was increased during activation of the mesoaccumbens and mesocortical pathways, whereas win-stay and safe-stay behavior were unaffected (Fig. 4d and Fig. S6c). Thus, activation of both ascending VTA projections made animals less prone to alter choice behavior after losses, which significantly impaired task performance during mesoaccumbens activation. The increase in lose-stay behavior during mesocortical activation is the result of the preference for the risky lever in the second trial block, but this did not result in poor choice behavior (Fig. 4c).

To test whether the effects in this task were specific to devaluation mechanisms, we trained the animals expressing DREADD in mesoaccumbens neurons on the same task with increasing, instead of decreasing odds of reward at the risky lever (Fig. 4e). In this condition, mesoaccumbens activation did not significantly change risky choice in any of the blocks (Fig. 4f), although a modest but significant decrease was observed in performance (i.e. a lower fraction of optimal choices; Fig. 4g) which was caused by a higher preference for the risky lever in the first few trials (Fig. S6d). This could be the result of a reduced ability of the animals to devalue the outcome of responding on the risky lever in the initial trials of the first block. However, since this version of the task primarily relies on revaluation, rather than devaluation mechanisms, especially in later blocks (Fig. S6b), a mesoaccumbens stimulation-induced devaluation deficit caused no further changes in behavior. Indeed, win-stay and lose-stay behavior were unaffected by mesoaccumbens activation (Fig. 4g).

In sum, the effects of chemogenetic activation on the probabilistic discounting task support our hypothesis that mesoaccumbens activation results in an inability of animals to adapt behavior to lower-than-expected outcomes, which under physiological circumstances is mediated by negative RPE signals in DA cells. In contrast, mesoaccumbens hyperactivity did not markedly interfere with adaptations to higher-than-expected outcomes. Furthermore, mesocortical activation increased risky choice behavior, but only when this was without negative consequences for the net gain in the task.

### Dopamine pathway activation does not change static reward value

Changes in static reward value may influence behavior in tasks investigating dynamic changes in reward value, such as the reversal learning task. For example, food rewards may be less or more appreciated due to changes in feelings of hunger, satiety or pleasure. Alternatively, operant responding may become habitual rather than goal-directed when manipulating the striatum, although this is thought to be mediated by its dorsal parts rather than the NAc^22,36^.

To assess whether alterations in static reward value or in the associative structure of operant responding contributed to the behavioral changes evoked by DA pathway stimulation, rats were subjected to operant sessions in which they could lever press for sucrose under an FR-10 schedule of reinforcement. Activation of the mesoaccumbens and mesocortical pathways did not alter the total number of lever presses (Fig. 5a), suggesting that absolute reward value was unchanged. We also tested animals in operant sessions, whereby in half of the sessions the animals were pre-fed with the to-be obtained reward. This type of devaluation tests whether animals retain the capacity to adjust operant behavior to changes in (the representation of) reward value. Pre-feeding robustly diminished lever pressing for sucrose, both in a non-reinforced extinction session, as well as under an FR5 schedule of reinforcement. Importantly, this effect of chronic devaluation was not affected by mesoaccumbens or mesocortical activation (Fig. 5b), indicating that responding remained goal-directed^36^.

**Figure 5.**
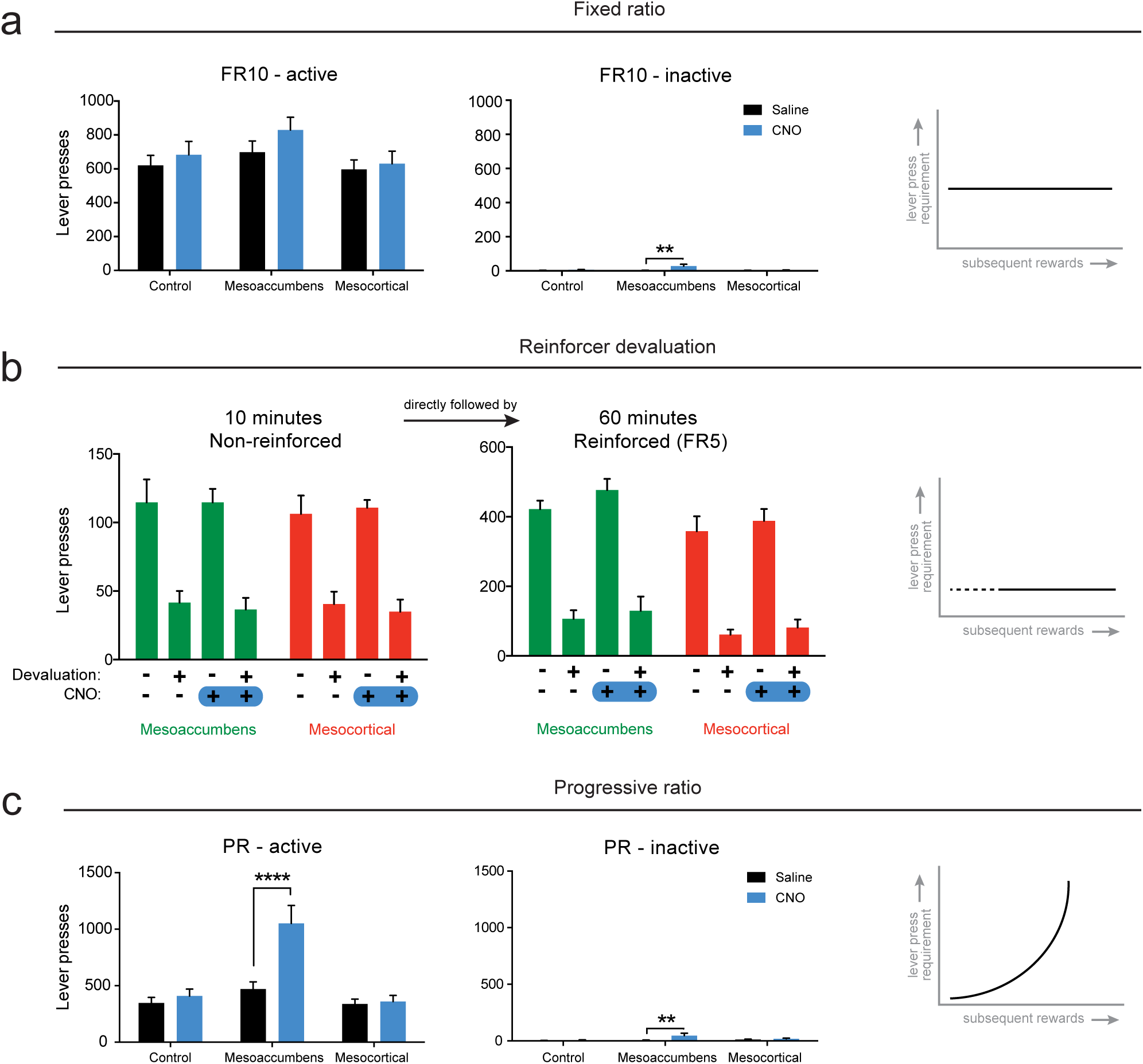
Mesocortical and mesoaccumbens activation does not alter the static reward value of sucrose. (**a**) DREADD activation of either pathway did not affect the number of active lever presses for sucrose under a fixed-ratio 10 schedule of reinforcement (two-way repeated measures ANOVA; main effect of CNO, p = 0.0355; group × CNO interaction, p = 0.5001; post-hoc Sidak’s multiple comparisons test, CNO versus saline, all p > 0.1). A significant but numerically modest increase was observed in inactive lever presses after mesoaccumbens activation (two-way repeated measures ANOVA; main effect of CNO, p = 0.0096; group × CNO interaction, p = 0.0207; post-hoc Sidak’s multiple comparisons test, CNO versus saline, p = 0.9302 for controls, p = 0.0017 for mesoaccumbens group; p = 0.9957 for mesocortical group). n = 9 for control, n = 8 for mesoaccumbens group, n = 9 for mesocortical group. (**b**) Both during a 10-minute extinction session (left panel) and a reinforced lever pressing session (under an FR5 schedule of reinforcement, right panel), devaluation of the reinforcer by selective satiation for sucrose lead to a decrease in responding (2-way repeated measures ANOVA, main effect of prefeeding in all four groups, p < 0.0001), without any effects of CNO (non-reinforced mesoaccumbens, CNO effect p = 0.7745, prefeeding × CNO interaction: p = 0.8448; non-reinforced, mesocortical, CNO effect p = 0.9516, prefeeding × CNO interaction: p = 0.5318; reinforced mesoaccumbens, CNO effect p = 0.1472, prefeeding × CNO interaction: p = 0.5287; reinforced mesocortical, CNO effect p = 0.4654, prefeeding × CNO interaction: p = 0.8877). n = 12 for mesoaccumbens, n = 11 for mesocortical group. (**c**) Under a progressive ratio schedule of reinforcement, mesoaccumbens activation significantly increased the number of lever presses made (two-way repeated measures ANOVA; main effect of CNO, p = 0.0006; group × CNO interaction, p = 0.0007; post-hoc Sidak’s multiple comparisons test, p = 0.8998 for controls; p = 0.8998 for control group; p < 0.0001 for mesoaccumbens group; p = 0.9947 for mesocortical group). A significant but numerically modest increase in cumulative inactive lever presses was observed after mesoaccumbens stimulation (two-way repeated measures ANOVA; main effect of CNO, p = 0.0204; group × CNO interaction effect, p = 0.0680; post-hoc Sidak’s multiple comparisons test, CNO versus saline, p = 0.9840 for controls; p = 0.0082 for mesoaccumbens group; p = 0.9392 for mesocortical group). n = 9 for control, n = 8 for mesoaccumbens group, n = 9 for mesocortical group. Data are shown as mean ± standard error of the mean.

Consistent with previous findings^37,38^, activation of the mesoaccumbens pathway increased operant responding under a progressive ratio schedule of reinforcement^39^ (Fig. 5c), which is often interpreted as reflecting an increased motivation to obtain food^37–39^. However, in light of the present findings, we interpret this finding to reflect that that mesoaccumbens activity renders animals less able to devalue the relative outcome of pressing the active lever when the response requirement increases over the session, hence leading to increased response levels. Such an action devaluation likely involves negative RPE signals from DA neurons.

### Mesoaccumbens hyperactivity evokes punishment insensitivity

To test whether the devaluation deficit as a result of mesoaccumbens hyperactivity also resulted in an inability to incorporate explicitly negative consequences into a decision, we subjected animals to a novel punishment task, in which reward taking was paired with an increasing chance of an inescapable footshock (Fig. 6a). As expected, the introduction of this 0.3 mA footshock punishment diminished responding for sucrose, an effect that persisted after injection of CNO in the mesocortical and sham control groups (Fig. 6b). In contrast, activation of the mesoaccumbens pathway completely abolished this punishment-induced reduction in responding, as the animals took as many rewards as under non-punishment conditions. This finding suggests that during mesoaccumbens hyperactivity, reward value is not properly discounted — in other words, animals are not able to take the increasingly negative consequences of an action into account. Consistent with a role for DA neurotransmission in processing these punishment signals, we observed, using *in vivo* calcium imaging, that footshock evoked a reduction in the activity of VTA DA neurons (Fig. 6c).

**Figure 6.**
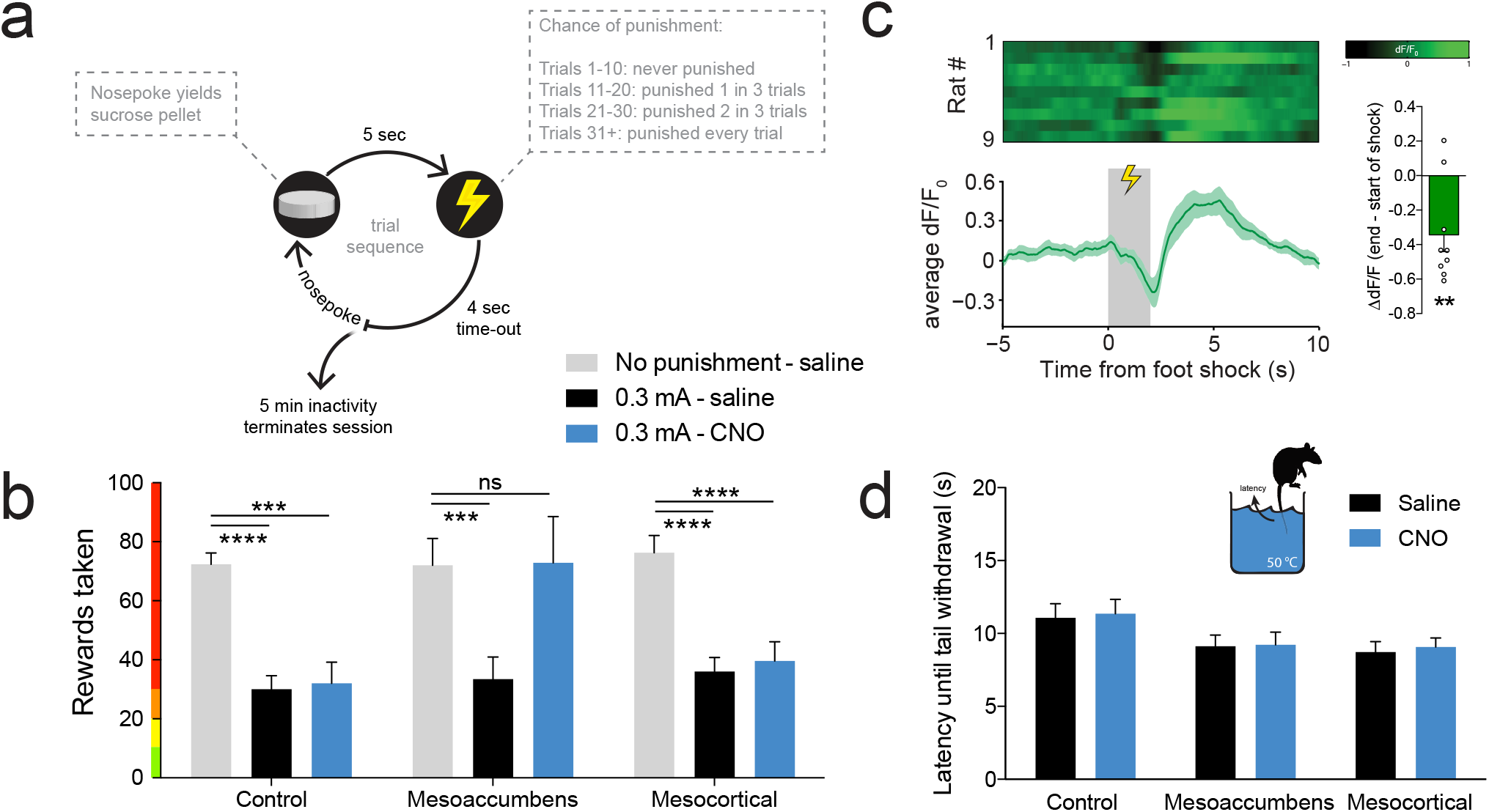
Mesoaccumbens, but not mesocortical activation attenuates the effect of punishment on responding for sucrose. (**a**) Task design. (**b**) After saline treatment, footshock punishment robustly diminished responding (Sidak’s multiple comparisons test, ‘0.3 mA saline’ versus ‘no punishment saline’, all p < 0.001). This effect was abolished by activation of the mesoaccumbens, but not the mesocortical, pathway (Sidak’s test, ‘0.3 mA CNO’ versus ‘no punishment saline’ in the mesoaccumbens group, p = 0.9995; in mesocortical group, p = 0.0002; in control group, p < 0.0001). n = 9 control, n = 9 mesoaccumbens group, n = 10 mesocortical group. (**c**) Footshock punishment evoked a decrease in DA neuron activity, measured using fiber photometry in TH::Cre rats (one-sample t-test, p = 0.0074, n = 9 rats). (**d**) No modulation of nociception by mesoaccumbens or mesocortical activation in the tail withdrawal test (2-way repeated measures ANOVA; main effect of CNO, p = 0.75; group × CNO interaction, p = 0.99). n = 8 control, n = 9 mesoaccumbens group, n = 9 mesocortical group. Data are shown as mean ± standard error of the mean. **** p < 0.0001, *** p < 0.001.

To control for effects on nociception in our punishment task, we subjected the animals to a tail withdrawal test, and found this not to be affected by mesoaccumbens activation (Fig. 6d). Moreover, anxiety, as tested in the elevated plus maze (Fig. S7a,b), was unaffected by mesoaccumbens stimulation. Consistent with literature, we found that mesoaccumbens stimulation increased locomotion (Fig. S8a), just like cocaine and D-amphetamine do^40,41^. We think, however, that the changes in value-based decision-making observed in the punishment task, as well as in the other tasks, cannot readily be attributed to increased locomotion. First, reaction times in the punishment task were longer after mesoaccumbens activation (Fig. S8b). Second, responding in the inactive hole in the punishment task was not changed (Fig. S8c). Third, the effects of mesoaccumbens activation in the reversal learning task were restricted to win-stay behavior after the first reversal. Last, mesoaccumbens activation did not affect the time for the animals to complete the reversal learning session (Fig. S3d).

### RPE processing during mesoaccumbens hyperactivity

There are three possible explanations for the impaired negative RPE processing during mesoaccumbens hyperactivity: (1) hyperactivity of VTA DA neurons abolishes the trough in neuronal activity caused by negative reward prediction, (2) elevated DA levels lead to a baseline shift in RPE signalling, after which a decrease in DA release during negative reward prediction does not reach the lower threshold necessary to provide a learning signal in downstream regions, or (3) a combination of both.

To address the first explanation, we unilaterally injected animals with a mixture of the calcium fluorophore GCaMP6s and Gq-DREADD and tested animals for reversal learning (Fig. 7a and Fig. S9). This allowed us to measure RPE signals from VTA neurons within one animal during baseline conditions and during hyperactivation of these same neurons. CNO administration did not impair the ability of VTA DA neurons to signal RPEs during reversal learning (i.e. deviations from baseline during reward prediction), inconsistent with the first possible explanation. By extension, this also excluded the third explanation. However, the second explanation is consistent with our findings that chemogenetic stimulation of the mesoaccumbens pathway increases the extracellular concentration of dopamine and its main metabolites in the NAc (Fig. 2h). Together, these data support a scenario in which the inability to adjust behavior after loss or punishment during hyperactivation of the mesoaccumbens pathway is not due to an inability of VTA neurons to decrease their firing rate during negative reward prediction, but rather by impaired processing of this learning signal within the NAc as a result of increased baseline DA levels (Fig. 7b). This observation fits well with our earlier finding that the infusion of a DA antagonist into the NAc can prevent the effects of DREADD activation on reversal learning (Fig. 2j), a manipulation that restores the degree of NAc DA receptor activation.

**Figure 7.**
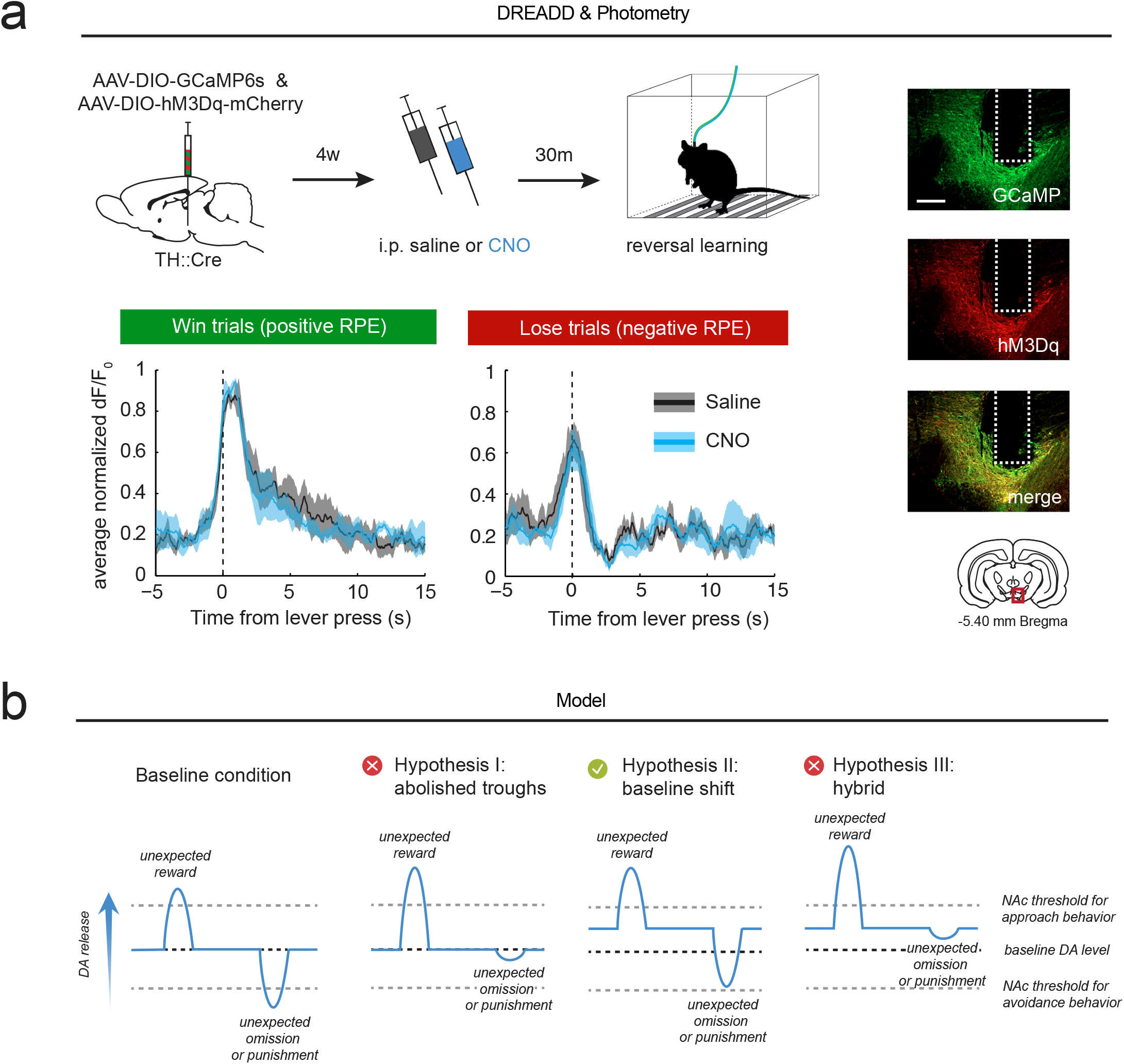
RPE processing after mesoaccumbens stimulation. (**a**) Animals were co-injected with GCaMP6s and Gq-DREADD and tested for reversal learning after injection of saline or CNO. VTA neurons responded in a comparable way during reversal learning after saline and CNO treatment (repeated measures in n = 4 animals; ANOVA, CNO x time interaction effect, win trials, p = 0.39; lose trials, p = 0.38). See figure S9a for individual animals. Scale bar, 1mm. Data are shown as mean (solid line) ± standard error of the mean (shading). (**b**) Proposed mechanisms: (I) Hyperactivity of NAc-projecting VTA DA neurons leads to impaired coding of negative RPE troughs, (II) Hyperactivity shifts baseline NAc DA levels, thereby preventing the exceedance of a negative RPE threshold in the NAc and impairing the ability to learn from negative feedback, or (III) A combination of both.

## DISCUSSION

Here, we show that hyperactivity of the mesoaccumbens pathway reduces the ability of animals to use loss and punishment signals to change behavior by interfering with negative RPE processing. Using *in vivo* neuronal population recordings, we show that the VTA signals reward presentation as well as reward omission during VTA neuron hyperactivity, meaning that the behavioral impairments are not caused by blunted DA neuron activity during negative reward prediction, but rather by impaired processing in the NAc as a result of elevated baseline levels of DA. Therefore, we propose a model (Fig. 7b) in which hyperactive VTA neurons signal positive and negative RPEs to the NAc, but because baseline DA tone is increased, the signaling threshold in the NAc that allows for the incorporation of negative RPEs into adaptive behavior cannot be reached during reward omission or punishment.

The majority of neurons transfected with the DREADD virus had a DAergic phenotype, chemogenetic mesoaccumbens activation replicated the effects of cocaine and D-amphetamine on reversal learning, and this effect of chemogenetic mesoaccumbens activation was prevented by intra-NAc infusion of the DA receptor antagonist α-flupenthixol. Together, this supports the notion that the behavioral changes observed in the present study are the result of chemogenetic stimulation of VTA DA cells. However, a role for non-DA neurons cannot be excluded with the currently used techniques. Importantly, alongside the dense DA innervation, the VTA sends GABAergic, glutamatergic, as well as mixed DA/GABA or DA/glutamate projections to the NAc and mPFC^16,42,43^. The role that these projections play in behavior is only beginning to be investigated, but on the basis of what is presently known, we consider it unlikely that the non-DAergic innervation of the NAc and mPFC is involved in the behavioral changes observed here. For example, optogenetic stimulation of VTA GABA neurons has been shown to suppress reward consumption, something we did not observe in our experiments^44^. In addition, by inhibiting NAc cholinergic interneurons, stimulation of VTA GABA projections to the NAc has been shown to enhance stimulus-outcome learning^45^. However, increased stimulus salience does not readily explain the deficits in reversal learning, probabilistic discounting and punished responding for sucrose that we found in the present study. Last, stimulation of VTA-NAc glutamate neurons has been shown to produce aversive effects^46^, which in our experiments most likely would have increased, rather than impaired the ability to use negative feedback to alter behavior. Therefore, we think it is justified to state that the deficits in reversal learning, probabilistic discounting and punished reward taking evoked by chemogenetic mesoaccumbens stimulation is the result of increased DA signaling in the NAc. Reversal learning impairments have previously been reported after systemic or intra-NAc treatment with a DA D2 receptor agonist in rats and humans^47–49^, whereas probabilistic discounting seems to be dependent on DA D_1_ rather than D_2_ receptor stimulation in the NAc^50^. Together, this suggests that the behavioral effects of mesoaccumbens hyperactivity observed here rely on stimulation of both DA receptor subtypes, depending on the task structure. Interestingly, the punishment insensitivity we observed after mesoaccumbens stimulation appears inconsistent with previous studies showing that treatment with amphetamine and the DA D_2_ receptor agonist bromocriptine make animals more sensitive to probabilistic punishment in a risky decision-making task, in which animals can choose between a small and safe reward, and a large reward with a chance of punishment^51,52^. In this latter task, however, presentation of the punishment coincides with the presentation of the large reward, and it is unknown how DA neurons respond to such an ambivalent combination of events. Importantly, risky choice behavior was found to correlate positively with DA D1 receptor expression in the NAc shell^52^, suggesting that the influence of NAc DA on behavior in this task may not be unidirectional.

In contrast to the mesoaccumbens projection, hyperactivity of the mesocortical pathway did not markedly affect value-based decision-making. It did increase the preference for large, risky rewards over small, but safe rewards in the probabilistic discounting task. However, when one of the two options yielded more sucrose reward, animals remained capable of choosing the most beneficial option, perhaps as a result of the differential roles that prefrontal D1 and D2 receptors play in this task^53^. That these animals maintained the capacity to make proper value-based decisions was also apparent in the reversal learning and punishment tasks. Thus, the patterns of effects of mesocortical stimulation is qualitatively different from the mesoaccumbens-activated phenotype, even though there is modest overlap, such as the increased lose-stay behavior in the probabilistic discounting task. Therefore, we do not think that the mesocortical phenotype is an attenuated version of the mesoaccumbens one, although the lower density of the mesocortical projection (Fig. S2a) may explain the relative paucity of behavioural changes after chemogenetic mesocortical stimulation. Notably, the mesocortical pathway has been shown to be vital for certain forms of cost-benefit judgement, especially those involving uncertainty or sudden changes in task strategy^25^. As a result, manipulations of prefrontal DA affect tasks like probabilistic discounting or set shifting, but not reversal learning^25,54^.

Our data emphasize the importance of balanced DA signaling in the NAc. It is reasonable to assume that brain DA concentrations are tuned to levels that are optimal to survival, and deviations from this optimum lead to the profound behavioral impairments seen in certain mental disorders. We think that our proposed model of mesoaccumbens overactivation can explain the decision-making deficits that are seen during states of increased DAergic tone, such as manic episodes, substance abuse, and DA replacement therapy in Parkinson’s disease. When one cannot devalue stimuli, actions or outcomes based on negative feedback, their value representation remains artificially elevated. Hence, outcome expectancies of choices will be unrealistically high, leading to behavior that is overconfident and overoptimistic. These inflated outcome expectancies have been demonstrated in human manic patients^2^, suggesting an inability to devalue goals towards realistic levels. That this disease state is associated with abolished negative RPE signaling in the NAc is substantiated by an fMRI study in patients experiencing acute mania^55^, in which activity in the NAc of manic patients remained high when monetary reward was omitted, while healthy controls showed a significant reduction in NAc activity, as expected based on RPE theory.

Most drugs of abuse enhance DA transmission in the brain, either in a direct (e.g. DA reuptake inhibition) or indirect way (e.g. disinhibition of DA neurons)^56,57^. Direct dopaminomimetics, such as cocaine and D-amphetamine, are known to mimic the symptoms of mania, such as increased arousal, euphoria, and a reduced decision-making capacity^10^. Impaired learning from negative feedback may potentially contribute to the escalation of drug use, since users may be insensitive to the thought of forthcoming negative consequences during the ‘high’ of these drugs. Furthermore, DA replacement therapy, often prescribed to Parkinson’s disease patients, has been associated with the development of problem gambling, hypersexuality and excessive shopping behavior, a phenomenon known as the DA dysregulation syndrome^58,59^. More than a decade ago, it has already been hypothesized that these clinical features could be the result of impaired RPE learning due to ‘overdosing’ midbrain DA levels^30,60^. Here, we provide direct evidence to support this notion.

## Conclusion

There is a wealth of evidence to implicate increased DA levels in harmful decision-making behavior in mental disorders^1,2,3^. Thus far, however, it was unknown through which pathways and by which mechanisms these effects were mediated. Here, we used behavioral tasks in rats, combined with projection-specific chemogenetics to show that hyperactivation of the VTA leads to decision-making deficits by impairing negative feedback learning through overstimulation of NAc DA receptors. Altogether, we provide a mechanistic understanding of why decision-making goes awry during states of hyperdopaminergic tone, providing an explanation for the reckless behaviors seen during drug use, mania, and DA replacement therapy in Parkinson’s disease.

## ACKNOWLEDGEMENTS

Clozapine-N-oxide was a generous gift from the NIMH Chemical Synthesis and Drug Supply Program. We thank Roshan Cools for giving feedback on the manuscript, and the entire Adan and Vanderschuren labs for helpful discussions and feedback. This work was supported by the European Union Seventh Framework Programme under grant agreement number 607310 (*Nudge-IT*), and the Netherlands Organisation for Scientific Research (NWO) under project numbers 912.14.093 (*Shining light on loss of control*) and 863.13.018 (*NWO/ALW Veni grant*).

## AUTHOR CONTRIBUTIONS

J.P.H.V., J.W.D.J., G.v.d.P., R.A.H.A. and L.J.M.J.V. designed the experiments. J.P.H.V., J.W.D.J., T.J.M.R., C.F.M.H., R.v.Z., M.C.M.L., G.v.d.P. and R.H. performed the experiments. J.P.H.V. analyzed the behavioral and calcium imaging data. J.P.H.V. performed and H.E.M.d.O. supervised the computational analysis. I.W. and R.H. analyzed the microdialysis experiments. J.P.H.V., H.E.M.d.O., R.A.H.A. and L.J.M.J.V. wrote the paper with input from the other authors.

